# The emergence of bacterial blight pathogen followed the dispersal pattern of rice in Asia

**DOI:** 10.64898/2026.02.19.706742

**Authors:** Ian Lorenzo Quibod, Marian Hanna Nguyen, Genelou Atienza-Grande, Sujin Patarapuwadol, Wichai Kositratana, Nafisah Nafisah, Celvia Rosa, Joko Prasetiyono, Fatimah Fatimah, Gouri Sankar Laha, Raman Meenakshi Sundaram, Alvaro L. Perez-Quintero, Dante Adorada, Gavin Ash, Casiana Vera Cruz, Van Schepler-Luu, Ricardo Oliva

**Affiliations:** International Rice Research Institute, Los Baños, Philippines; Centre for Crop Health, Institute for Life Sciences and the Environment, University of Southern Queensland, Queensland 4350, Australia; Department of Plant Pathology, Faculty of Agriculture at Kamphaeng Saen, Kasetsart University, Nakhon Pathom, Thailand; Indonesian Center for Rice Research, West Java, Indonesia; Research Center for Genetic Engineering, National Research and Innovation Agency, Indonesia, Bogor, Indonesia; ICAR-Indian Institute of Rice Research, Rajendranagar, Hyderabad, Telangana State, India; Plant Health Institute of Montpellier (PHIM), Université Montpellier, IRD, CIRAD, INRAE, Institut Agro, France; The Alliance of Bioversity International and CIAT, Cali-Palmira 763537, Colombia

**Author notes:** corresponding authors: Ricardo Oliva, Van Schepler-Luu.

## Abstract

Crop domestication has a significant effect on the evolutionary trajectory of plant pathogens by providing new ecological niches and abundant resources. The domestication of Asian rice (*Oryza sativa*) in Asia, approximately 9,000 years ago, might have shaped the genetic makeup of associated microbes into modern threats. In this study, we provide insight into the evolutionary history and dispersal pattern of the rice bacterial blight (BB) pathogen, *Xanthomonas oryzae* pv. *oryzae* (Xoo), one of the most destructive rice pathogens in the last century. The analysis of 433 Asian Xoo (AXoo) genomes identified twelve modern populations derived from three ancestral lineages (AXooL). Each population emerged with a unique genetic composition including the combination of pathogenicity factors. Bayesian reconstruction suggests that Xoo lineages emerged alongside *O. sativa* domestication hotspots and followed the dispersal pattern of rice across the continent. An ancient Xoo lineage (AXooL1) emerged in China and was likely dispersed with *japonica* rice. A second lineage (AXooL2) which could have turned up from China and spread across India, then evolved due to the domestication and spread of *indica* rice, and later on expanded eastward of Asia.. We also showed that recombination played a significant role in the emergence of AXooL3, which appeared more recently and might have spread with the rice trading routes. Our study aligns the evolution and dissemination of the phylogroup AXoo with the history of *O. sativa*, offering valuable insights for the formulation of precise disease management strategies.

**AUTHOR SUMMARY:** Rice domestication was a crucial step in the development of Asian civilization. However, this process also affected the evolution of an associated pathogen, leading to its emergence as a global threat. Rice bacterial blight (BB), caused by the pathogen *Xanthomonas oryzae* pv. *oryzae* (Xoo), has been a scourge in many Asian countries. Using population genomics, we explored the diversity and evolutionary history of Xoo in Asia (AXoo). Here we show that two ancestral pathogen lineages emerged in rice domestication centers (China and India) and dispersed with rice across the continent. More recently, recombination played a crucial role in the appearance of a third lineage that spread through trading activity. This study provides the implications of the adaptation of AXoo in *Oryza sativa,* and might be valuable in forecasting BB outbreaks.

## INTRODUCTION

Understanding the origin of crop pathogens is often possible by tracing the domestication pattern of their host plants. Studying those centers of domestication provides valuable insights into the coevolution of both organisms, enabling the identification of resistance factors or the isolation of virulent components. The domestication of crops such as maize, wheat, and potatoes has been linked to the emergence and dispersal of pathogens [1–4]. Significant efforts have been made to leverage the genetic diversity unfolded in these regions for crop protection purposes [5,6].

From a historical perspective, the domestication and cultivation of rice in Asia (*Oryza sativa*) allowed unique socioeconomic and demographic changes that shaped the Asian civilization [7]. Currently, paddy rice ranks among the top ten most produced agricultural commodities globally, with Asia contributing to 90.5% of the total production [8]. The domestication histories of the two major subspecies of Asian rice, *indica* and *japonica*, have been widely debated among researchers. One theory is that both were domesticated independently in different regions of Asia [9,10] while another suggests that *japonica* was first domesticated in China, and later transferred domestication-linked alleles to a proto-*indica* population in South Asia [11]. However, it is possible that the domestication process was much more complex, involving multiple origins and potential hybridization events with wild relatives [12,13]. Subsequent diversification and dispersal of rice were likely influenced by other factors, such as global temperature changes and human activities [14].

Bacterial blight (BB) is a major rice disease caused by *Xanthomonas oryzae* pv. *oryzae* (Xoo), and was first discovered in Japan in 1884 [15]. Since then, it has remained a significant issue in tropical and subtropical rice-growing regions worldwide. The pathogen’s biology, ecology, and distribution have been extensively described [16]. Xoo is a vascular pathogen that gains entry through natural leaf openings or wounds and uses various virulence factors to colonize rice tissues and suppress host defense responses [16]. This pathogen has evolved into diverse pathotypes that can infect different rice cultivars, each one carrying a diverse set of virulent factors [17–20]. However, the origin and emergence of Xoo are still unclear, and less information is available about the impact of rice domestication and dispersal on the evolutionary history of the pathogen. Through a detailed analysis of genomic datasets, this study aims to uncover the evolutionary process of Xoo and its connection to the domestication and dispersal of rice in Asia.

## RESULTS AND DISCUSSION

### Modern Asian Xoo populations emerged from three ancestral lineages

To characterize the genetic ancestry of Xoo in Asia (AXoo), 433 genomes were gathered from various sources (S1 Data). This collection spans over 50 years of bacterial blight in rice-growing regions in China (179), India (102), the Philippines (96), Thailand (32), Indonesia (12), Japan (5), South Korea (3), Taiwan (2), Nepal (1), and Australia (1). A maximum likelihood phylogenetic analysis of 22,115 non-recombining SNPs revealed three ancestral lineages named AXooL1, AXooL2, and AXooL3 (Fig 1a). The same result was also observed when using Xoc as an outgroup (S1 Fig). Previous studies have also recognized three major Xoo lineages in East Asia, South Asia, and Southeast Asia using different genetic markers [17,21,22], which supports our result. A population structure analysis using the BAPS (Bayesian Analysis of Population Structure) approach [23] further classified the three lineages into 12 modern populations: AXoo1-AXoo12 (Fig 1a; S2a Fig). This clustering was also supported by the evolutionary distance estimation and pairwise SNP distance matrix (S1b-c Fig). Furthermore, the population structure from a previous analysis from China, India, and the Philippines [19,20,22] coincided with the predicted groupings resulting from this analysis (S2a Fig).

**Fig. 1.**
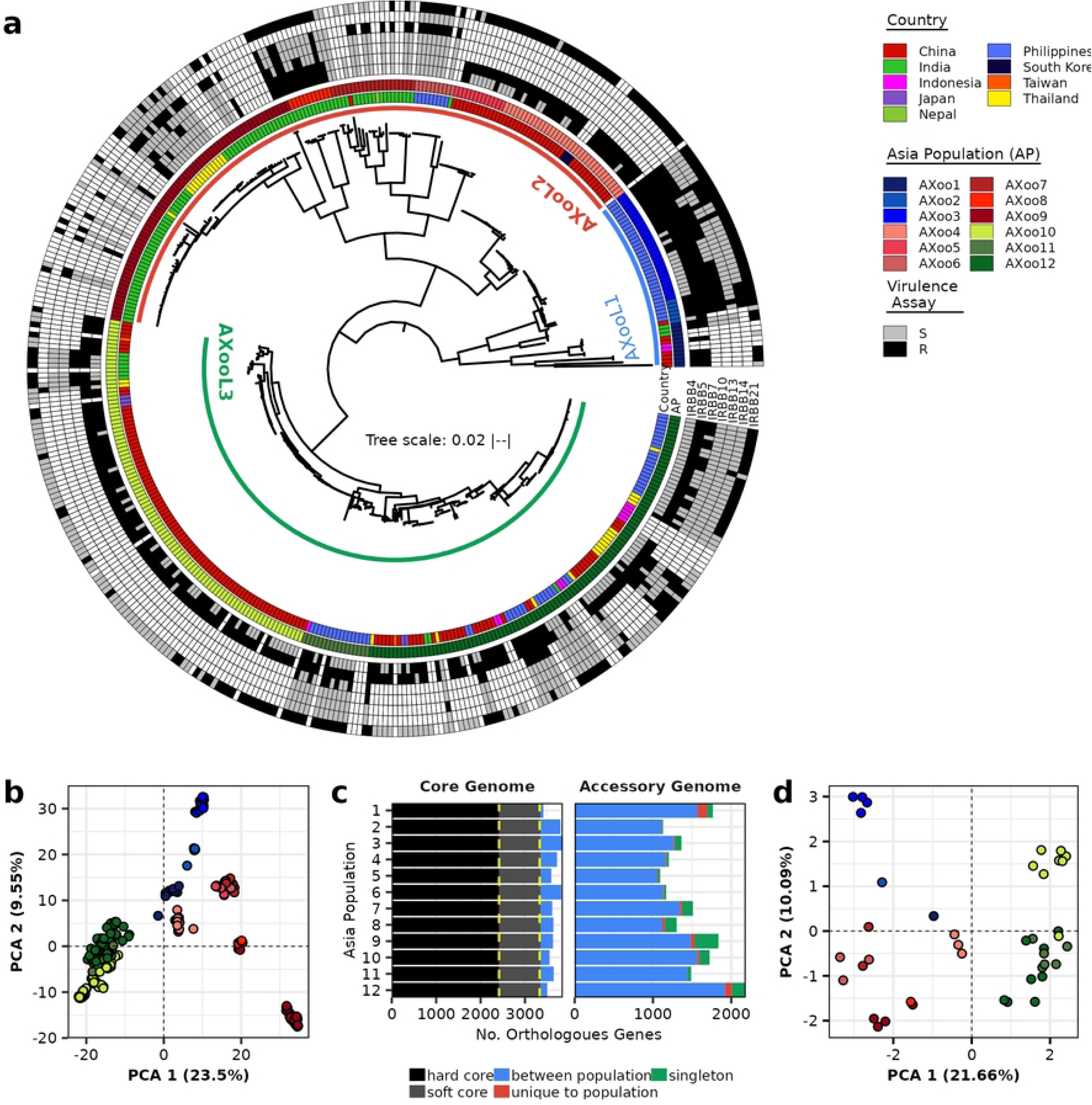
Phylogenetic relationship, population structure, and pan-genome of *Xanthomonas oryzae* pv. *oryzae* strains collected in Asia between 1965 to 2018. a) A maximum-likelihood tree was constructed using 22,115 non-recombining genome-wide SNPs. The Asian Xoo (AXoo) isolates were grouped into three lineages (AXooL1 to AXooL3) and twelve modern Asian populations (AXoo1 to AXoo12) which were inferred using fastbaps [23]. The inner ring corresponds to the isolate’s country of origin. The second ring indicates the population predicted by the fastbaps analysis. The other seven rings show virulence reactions (S: susceptible, R: resistant) to a set of near-isogenic lines carrying single resistance genes (*Xa*). White filled lines denote no data available. b) Principal component analysis (PCA) of the 7,585 orthologous gene alleles identified in the pan-genome. c) The orthologues gene content of each population and its distribution as part of the core or accessory genome. The pan-genome was partitioned as hardcore (present in all isolates), softcore (present in 95% of the isolates), between-populations (present in some of the populations), unique to population (gene group present only in one population), and singleton (present only in a single isolate). The orthologous genes belonging to the between-population classification can contain genes in the core and accessory genomes. The yellow dotted vertical lines serve as boundaries. d) Principal component analysis using allelic variations of 34 Transcription activators-like (TAL) effectors and 26 Xanthomonas outer proteins (Xop) effectors identified from completely assembled genomes.

The AXoo isolates were collected from major river basins, islands, and archipelagos in South Asia, East Asia, mainland Southeast Asia, and archipelagic Southeast Asia (Fig 2a). There seems to be a pattern of regional dispersal from this collection. For instance, AXoo5, AXoo9, and AXoo7 populations were found in mainland Asia while AXoo3 and AXoo11 populations were observed in the Southeast Asian archipelagos. Other populations, such as AXoo1, AXoo10, and AXoo12, have dispersed regionally across rice-growing areas. However, this pattern of distribution may change when more genomes are available.

**Fig. 2.**
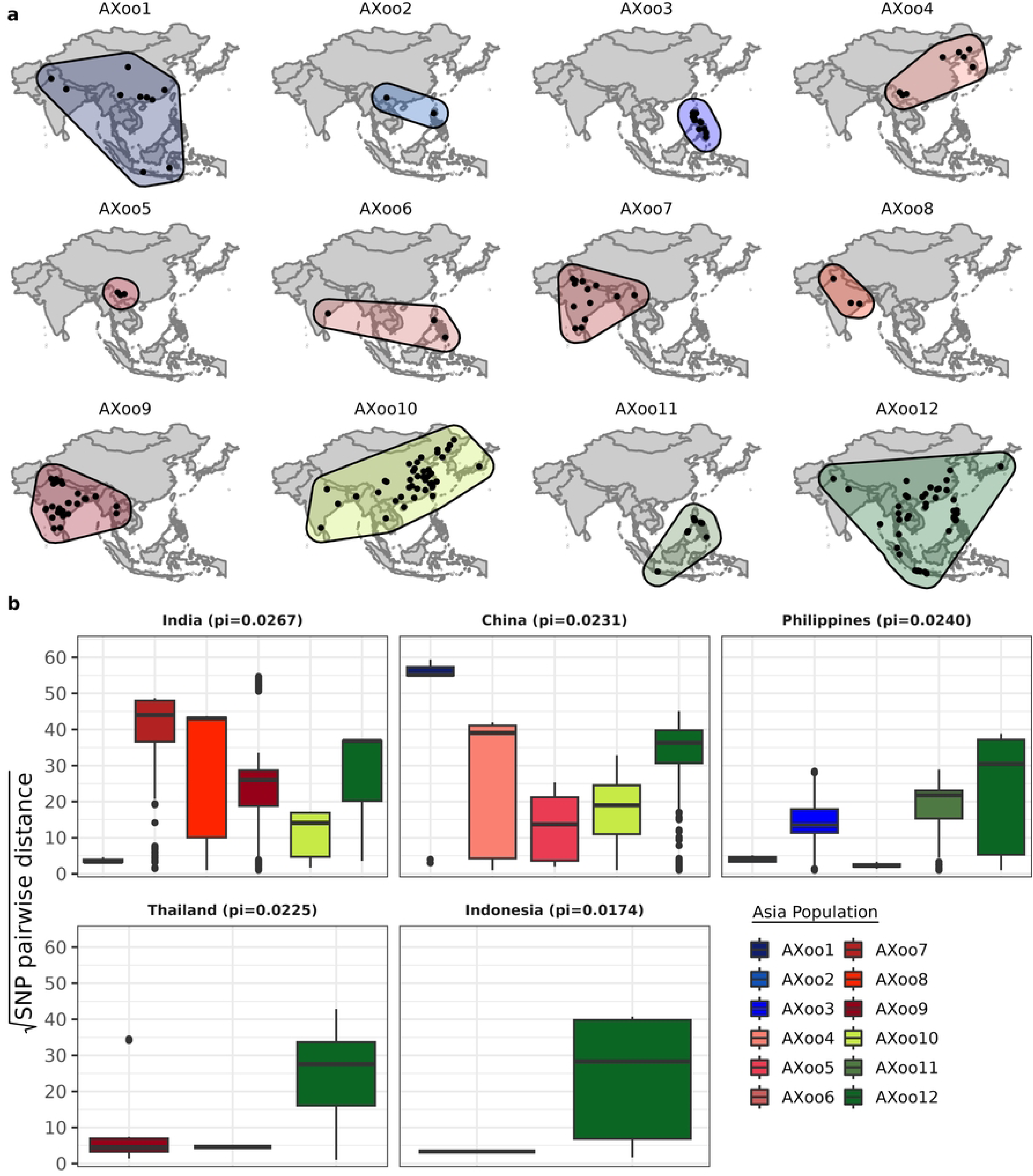
Distribution and diversity of *Xanthomonas oryzae* pv. *oryzae* populations in Asia (AXoo). a) Geographical landscape of 12 AXoo populations in Asia. The dots indicate the location of the isolates. Blue, red, and green shades represent AXooL lineages. b) Diversity distribution of AXoo in major rice-producing countries using pairwise SNP difference for each isolate. Nucleotide diversity (Pi) was computed for well-sampled countries with more than 10 genomes.

The center of origin of AXoo appears to be in southern China, especially in the Yunnan Province, given the extensive diversity observed in AXooL1 (AXoo1, AXoo2), AXooL2 (AXoo4, AXoo5, AXoo7), and AXooL3 (AXoo10, AXoo1). This is also the region with multiple sub-lineages circulating from each major lineage. While we cannot explain all evolutionary events occurring in each population due to sample limitation, some could be deduced. For instance, AXoo10 and AXoo12 have undergone rapid diversification and expansion across various Asian regions. One or two early westward dissemination events to India might be sufficient to explain the diversification of AXooL2 in India, and one of these populations then underwent recent clonal expansion (AXoo9).

Although other genetic groups may be absent from this collection due to the limited number of genomes from other countries, the collection of 433 isolates from the 12 AXoo populations, with spatiotemporal information, (Fig 2a; S3 Fig) allows for further analysis of the genetic composition, origin, and dispersal of Xoo in Asia.

### Asian Xoo populations have discrete genetic compositions

To understand the genetic composition of AXoo populations, we investigated their gene content, allelic variants, and selection signatures. The pan-genome structure was identified by classifying all the 1,566,958 predicted open reading frames into 7,585 orthologous genes comprising 3,352 core and 4,233 accessory genes (S4a Fig). A principal component analysis using the allelic variation of the orthologous genes resembled the 12 groups, which further supports the classification of 12 AXoo populations (Fig 1b). Accessory genes detected in all members of the same population reached up to 14.86% (AXoo3) (Fig 1c). Additionally, accessory genes belonging to a single population or a single strain reach up to 0.05% (AXoo1) or 0.08% (AXoo9), respectively (Fig 1c). Pairwise comparisons of the allelic genes belonging to the core (S4c Fig) and accessory (S4d Fig) groups showed that most AXoo populations within a lineage tend to have similar sets of alleles. AXoo12 shares the highest gene variants between populations with an average of 3,801 genes, while AXoo5 has the fewest number of shared genes (2,386 genes, Fig 1c), suggesting a possible difference in recombination ability between the two populations.

To estimate whether the 433 genomes would be sufficient to capture most orthologous gene repertoire of AXoo, the pan-genome was evaluated to see if it was open or closed. The power law model [24] showed that the AXoo has a closed pan-genome (α= 1.126 +/- 0.019, κ =654.466 +/-18.098) despite a large pan-genome repertoire, suggesting that adding more genomes would only yield a few new orthologous genes. Similar closed pan-genome patterns have been described in other plant pathogens, such as *Xanthomonas citri* pv. *citri*, the causal agent of citrus canker [25]. Similarly, the rarefaction curve for each AXoo population also predicted that adding more genomes would only yield a few new orthologue gene groups (S4b Fig). The only exception is population AXoo9 (α= 0.940 +/- 0.013, κ = 307.562 +/- 6.520), which also has the highest number of singletons (Fig 1c). Hence, more genomes for AXoo9 are needed to better capture the total number of orthologs. AXoo9 has been recently identified as the dominant group, known as L-1, in the rice-growing areas of India (S2a Fig) [19] and as a new virulent genotype overcoming *xa5* resistance gene in Thailand [26]. AXoo9-like strains have also been reported in Pakistan [27,28].

To assess genetic diversity among AXoo populations, nucleotide diversity (pi), genome-wide Tajima’s D (D) values, Fixation index (F_ST_), and the pairwise SNP number per isolate across regions were analyzed. AXoo1 was found to be the most diverse group with a two-fold higher than the average diversity index of all AXoo (Pi mean = 0.00026) (S5a Fig). AXoo7 (Pi mean = 0.00018) and AXoo12 (Pi mean = 0.00014) also showed high levels of diversity. The Tajima’s D values, ranging from −1.24 to 1.30, were significantly different from the null expectation suggesting a range of non-random events driving the population structure of Xoo in Asia (S5b Fig). For instance, the negative values found in AXoo3 (D median = −1.15), AXoo6 (D median= − 1.11), or AXoo11 (D median = −1.16), previously described in Quibod et al. [22], and might indicate recent population expansion or positive selection to remove variation. The population with the highest Tajima’s D positive value was AXoo8 (D median = 1.19) which likely reflects a drop in population size or the effect of balancing selection. The fixation index (F_ST_) among AXoo populations ranged from 0.19 to 0.72 (S5c Fig). AXoo1 showed, on average, the lower F_ST_ between populations (mean range = 0.34 to 0.54), indicating that it shares more genetic variation with most of the groups. In contrast, AXooL3 (F_ST_ mean range = 0.19 - 0.22) shared more genetic diversity within populations.

We then compared the distribution of the diversity index among the major rice-growing areas (Fig 2b). Our results showed that nearly 60% of the population are in China and India (Fig 2b). The AXooL1 in China had the highest median pairwise SNP number per isolate (pairwise SNP distance = 3,180) followed by the AXooL2 in India (pairwise SNP distance = 2,555) (Fig 2b). The AXooL3 diversity appears to be similar in all the regions. Although AXooL1 has the highest diversity in China, this lineage is infrequently sampled and more are typically found in South China, with the exception of some Northeastern China and Korean isolates. AXooL2 diversity primarily extends across South China, India and Thailand. While the AXooL3 is most abundant and rapidly diversifying across much of China and Southeast Asia. The mountainous geography of South China likely presented barriers to rice cultivation and pathogen dissemination for a long time after the divergence of AXooL1, L2 and L3. While the distribution of diversity in AXoo can be explained by population structure, the diversity and geographic distribution of AXoo populations may shift as more samples are collected. In areas with abundant sampling, AXoo lineages showed variation between China and India in the presence of different populations. Our data suggest a discrete evolutionary history of each AXoo population in Asia, where groups evolved distinct genomic features, and might have emerged within distinct geographies.

### Asian Xoo populations evolved unique phenotypes

To investigate the pathogenicity variation of AXoo populations, we first analyzed the reported phenotypic reactions of 344 Xoo strains over a set of seven near-isogenic lines (NILs) carrying single resistance genes (Fig 1a). The results suggest that members of the same AXoo population tend to exhibit similar phenotypic responses despite the variation in some NILs reactions. For instance, 80% of AXoo3 members cannot overcome the resistance gene *xa5* or *Xa21*. The same is true for AXoo4 members, which were unable to colonize plants with *xa5* or *Xa14*. Populations within AXooL3 showed the most variation, especially in response to *Xa4*, *Xa7*, and *Xa21* resistance genes. This highlights the need for proper resistance gene deployment across the Asian region as AXoo populations have varying capabilities to overcome resistance.

We also investigated the distribution and composition of type III effector genes across the AXoo population. Of the 433 genomes, only 41 genomes were assembled with long reads and could be used for annotating transcription activator-like effectors (TALEs). No TALE was annotated for AXoo5 due to the lack of long-read assembled genomes. We identified and classified the 693 TAL effectors from 41 strains (S2 Dataset) into 33 TALE Families (TEF) and 54 TALE clusters predicted by AnnoTALE (S6a Fig). Allelic variations in 33 TALE effectors and 26 *Xop* (Xanthomonas outer proteins) coding genes suggest that members of the same population seem to have similar effectors, which are occasionally shared with other populations across lineages (Fig 1d).

All AXoo populations have a minimum of one TALE allelic variant per locus (S6b Fig), and AXooL3 seems to carry more alleles than the other lineages. However, this could likely be due to the sampling bias since 6 to 8 genomes were available for the AXooL3 while others have only 1 to 4 genomes. Nevertheless, five TEFs, including AvrXa23, PthXo6, TEF19, TEF7/TEF26, and TEF9/TEF27, were present in all AXoo populations (S6b Fig) and may therefore be ancestral and have essential roles for the pathogen. While PthXo6 contributes to virulence by binding to the promoter of the bZIP transcription factor OsTFX1 [29], the virulence function of AvrXa23 remains unknown. Interestingly, AvrXa23 has high diversity in AXoo and is detected by a cryptic binding site on the cognate executor rice gene *Xa23*, which encodes a transmembrane protein from the wild species *O. rufipogon* [30]. A recent report has indicated that *Xa23* emerged within *O. rufipogon* populations in the Guangxi province [31] suggesting that a Xoo ancestor carrying *AvrXa23* might have evolved in that region over a considerable amount of time. Huang et al. [11] also pointed out that an *O. rufipogon* population from a similar region in South China was one of the ancestors of cultivated *japonica* rice.

TAL effectors *PthXo1*, *PthXo2*, *PthXo3*, and *AvrXa7* activate different sugar transporters from the *OsSweet* family [32,33]. All AXoo populations carry at least one *OsSweet*-inducing TALE, suggesting that sugar acquisition is an essential function for AXoo. Interestingly, AXoo12 carried all the *OsSweet*-inducing TALEs, including allelic versions of the *PthXo2* gene which can induce the expression of *indica* and the *japonica OsSweet13* [33]. Other TAL families, such as TEF19 and AvrXa10 were detected in a few populations, suggesting a derived origin. For example, AvrXa10, which activates the cognate executor resistance gene *Xa10* [34,35], was detected in populations AXoo2 and AXoo3. A recent study showed that AvrXa10 triggers nonhost resistance by targeting the promoters of C2H2-type zinc-finger protein orthologs in tobacco and tomato [36], and might play a critical role in targeting similar genes in other pathosystems. Overall, it seems likely that independent AXoo groups emerged and acquired unique fingerprints distinguishable by their effector repertoires and virulence profiles. Whether these features were shaped by host adaptation or other environmental factors remains unclear. However, the interpretations of effector presence across AXoo populations here may change as additional long-read genomes are incorporated since TALEs cannot be well resolved by short-read sequencing due to its highly repetitive nature.

### Recombination played a role in the emergence of lineage 3 populations

Recombination plays a key role in the evolution of plant pathogens [37,38]. Previous studies have detected discrete events of recombination across AXoo genomes [19,22,39]. To understand the overall pattern of recombination among AXoo lineages and populations, we characterize these events using multiple tools [40–42]. Inferred neighbor net analysis using 22,115 SNPs suggests that AXooL3 was highly affected by recombination, as evidenced by multiple reticulations in the SplitsTree4 network (Fig 3a), and significant PHI values (p-value = 0.00). Using the program ClonalFrameML, we estimated the overall recombination rate of AXoo. The analysis showed a ratio of recombination to mutation (R/θ) of 0.895, a mean DNA import length (δ) of 896 bp, and the divergence of each fragment by recombination (ν) at 0.00550. Unlike previous results, which showed that single nucleotide variations in AXoo are more likely to be introduced by point mutation than recombination [19,22], a lower value was observed at only 1.12 point mutations. The estimated ratio of single nucleotide substitutions introduced by recombination vs point mutation (r/m) was at 4.41, twice the earlier reported values of 2.23 [19] and 2.82 [22]. This discrepancy may be due to the fact that both previous studies only considered the effect of recombination in a single geographical origin, and our current data strongly support that AXoo are highly recombining. In contrast, a recent study showed r/m value for *Xanthomonas oryzae* species amounted to only 0.68 [43]. This suggests that recombination may occur more frequently within pathovars than between them.

**Fig 3.**
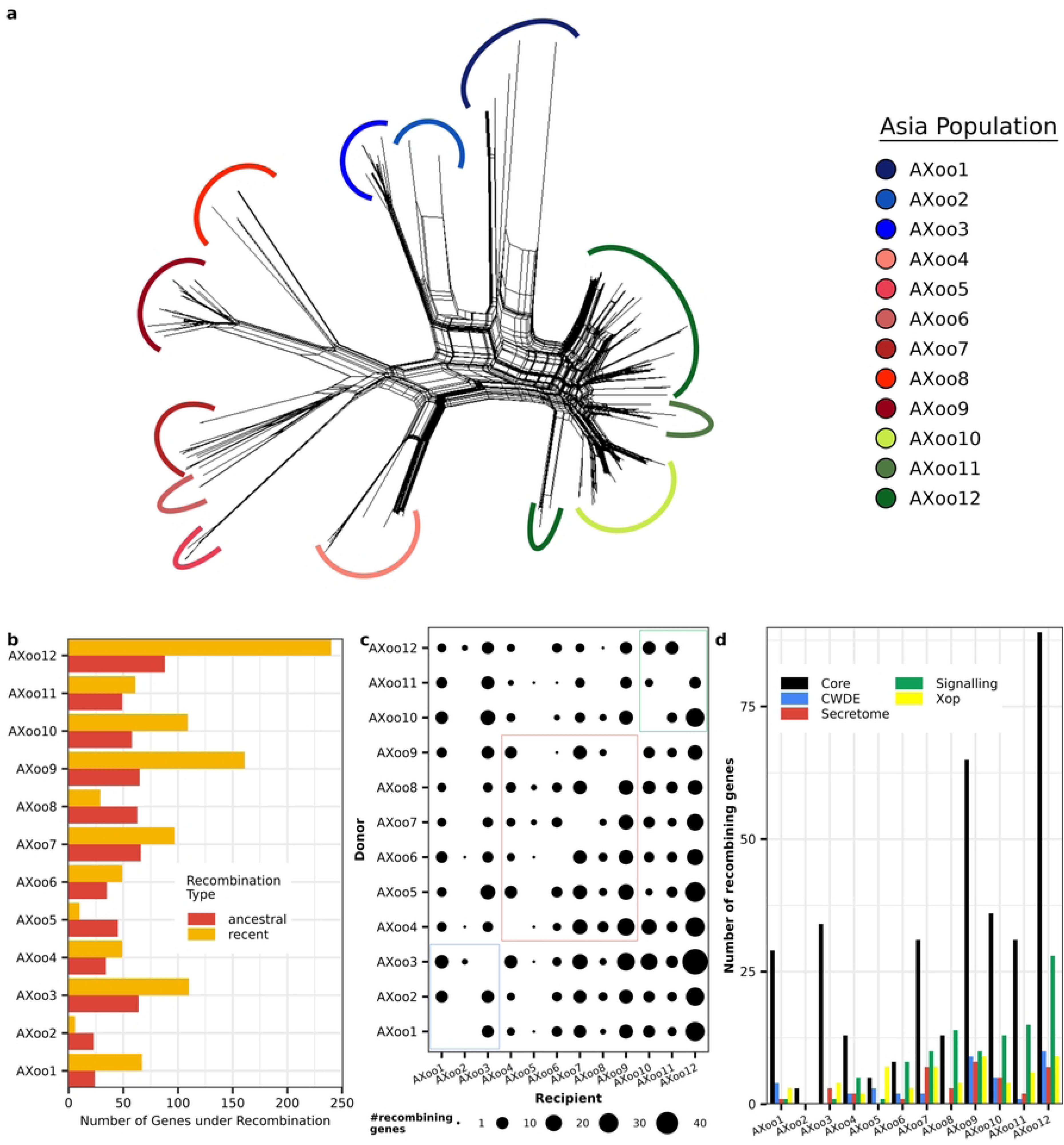
The recombination landscape of *Xanthomonas oryzae* pv. *oryzae* populations in Asia (AXoo). a) Phylogenetic network analysis of Xoo isolates using the neighbor net method implemented in SplitsTree4, where the twelve inferred AXoo populations are highlighted. b) A bar plot showing the distribution of genes under recombination per AXoo populations. The recombining genes were identified as ancestral (recombination affecting entire lineages) and recent (recombination affecting a few strains). c) A plot showing the flow of genes under recombination from the donor to the recipient AXoo population. The recent recombination was the only data used here. The size of the circle shows the total number of recombining genes between AXoo populations. d) A barplot showing the recombining gene distribution of the core and pathogenicity-related genes in each AXoo population. The colors reflect the different recombining gene classification: core (black), cell wall degrading enzyme/CWDE (blue), putative secreted gene/secretome (red), signaling and regulation of virulence/signaling (green), and the Xanthomonas outer proteins/*Xop* (yellow).

Furthermore, we used the orthologous genes to infer which of the populations have the strongest affinity for recombination (Fig 3b). The results of ancestral recombination (recombination involving the whole population) and recent recombination (recombination involving some of the isolates in each population), showed a significant impact on gene flow in AXoo12 with 328 recombinant genes (Fig 3b). Interestingly, most of the recombinant genes in AXoo12 were of the recent type, potentially representing a foundational event and responsible for the strong signal observed within AXooL3. AXoo9 was the second most recombinant population, with 226 recombinant genes, and AXoo6 (29 genes) was the least recombinant (Fig 3b). Regarding ancestral recombination, AXoo1 and AXoo2 of AXooL1 showed the lowest number of recombinant gene frequencies at 23 and 24 genes, respectively (Fig 3b). This might indicate that AXooL1 dispersal happened early, preceding any recombination events. Again, AXoo2 has the lowest number of recent recombinations, possibly due to the number of strains in this population.

Next, we assessed the frequency of recombinant genes in donor-recipient population pairs. The results suggested that AXoo populations are more likely to receive genes from members of the same lineage (Fig 3c). For instance, AXoo9 received approximately 18 genes from each donor population belonging to the same lineage. The same pattern was observed in other populations including AXoo1 and AXoo7. These results demonstrate that intra-lineage recombination was much more common compared to inter-lineage. The same observation was reported in the *P. syringae* species complex, where inter-phylogroup genetic exchange rarely occurs for the preservation of genetic cohesion [44]. In contrast, AXoo12 has donors coming from all the populations and received an average of 25 genes per donor population(Fig 3c). Hence, AXoo12 might have the potential to produce new virulent combinations [37,45]. To assess this feature, we inspected the recombination pattern in genes related to fitness and compared them with those of the core genes. As expected, pathogenicity-related genes such as cell-wall degrading enzymes (CWDE), signaling-secretome-related, and *Xop* genes were enriched in the recombination pattern of AXoo12 (Fig 3d). Previous studies have demonstrated that type III effectors from *Xanthomonas* species can be acquired through recombination [39,46], and play a significant role in the fitness and plasticity of pathogens. It is likely that one or several events of recombination have shaped the AXooL3 lineage, causing the appearance of new groups (AXoo12) in the rice ecosystem. Additionally, the overlapping distribution of multiple populations in one region (Fig 2a) provides more opportunities for recombination to introduce variation, as observed in AXoo12, for example. A recent study showed that the genomes of *Xanthomonas oryzae* are enriched with insertion sequence elements (IS) [47], and thus could also be a key to acquire new modifications through recombination. Overall, these findings clearly validate that recombination has a huge impact in the evolution of AXoo.

### Asian Xoo lineages could have emerged along with rice domestication

To estimate the divergence time of the ancestral Xoo groups, we used a Bayesian reconstruction approach [48] and found that AXoo lineages likely emerged at different time frames (Fig 4a). First, we performed a root-to-tip regression analysis, which showed a positive correlation in nine out of twelve AXoo populations (S7 Fig), signifying sufficient temporal signal. The smaller sample size and narrow temporal range in the remaining populations might explain the absence of an indirect correlation. To estimate the evolutionary rate of each AXoo population, all recombinant SNPs were excluded. The substitution rate per site per year in AXoo groups ranged from 8.63×10^-6^ in AXoo8 to 6.1810^-9^ in AXoo5-7, and there were different evolutionary rates within AXoo populations (S1 Table; S8a-e Fig). The nucleotide substitution rate coincided with the predicted genome-scale evolutionary rates in bacterial pathogens, which ranges from approximately 10^−5^ to 10^−8^ [49]. Furthermore, the observed substitution rates were comparable with other bacterial plant pathogens such as *Xanthomonas citri* pv. *citri* (8.4×10^−8^ and 8.2×10^-8^) [50,51], *Xanthomonas vasicola* (3.17×10^−7^)[52], and *Xylella fastidiosa* subspecies (whole species = 7.6×10^−7^, subsp. *fastidiosa* = 7.7×10^−7^, subsp. *multiplex* = 3.2×10^-7^) [53–55].

**Fig. 4.**
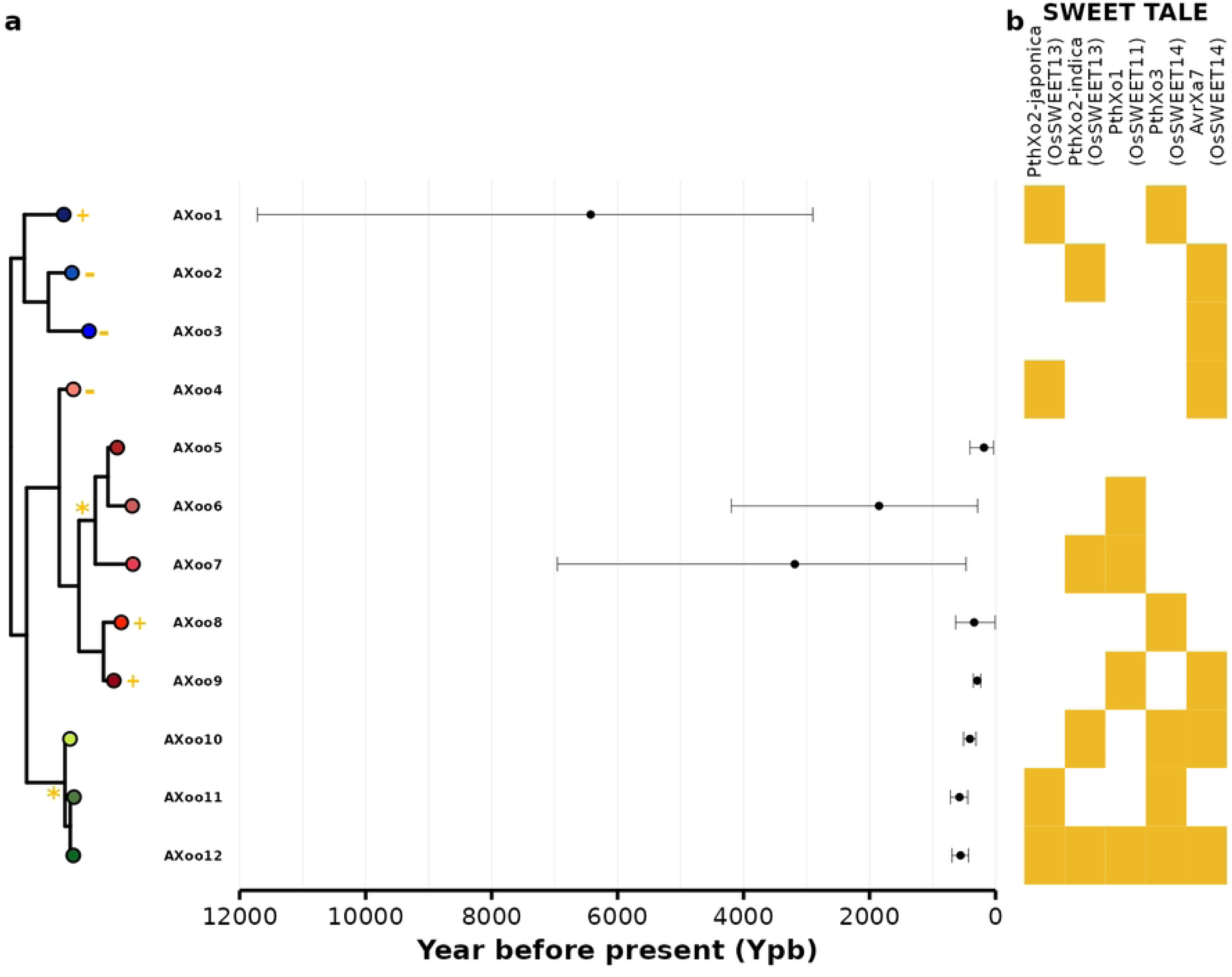
Emergence of *Xanthomonas oryzae* pv. *oryzae* populations in Asia and distribution of TALE targeting SWEETs. a) The graph shows the estimated timeline when the most recent common ancestor of each population emerged. The mean time is represented by a black dot with the intervals as the 95% highest posterior density (95% HPD) intervals. Some populations don’t have an estimated time of emergence because it failed to pass the initial testing for clock signals (Fig. S6). The yellow-orange symbols in the phylogeny signify which datasets have a temporal signal, and how the samples were tip-dated: + = positive temporal signal, but can only be analyzed individually, * = positive temporal signal and is analyzed by combining the population sharing the same node, and - = populations that have a negative temporal signal. The phylogeny is based on the collapse cluster of Fig 1. b) The heat map shows the distribution of TALEs targeting promoters of SWEET genes in *Oryza sativa*. The SWEET gene targets are shown inside the parenthesis. No long read sequences were available for strains belonging to the AXoo5 population.

Tip-dating analysis revealed the sequential emergence of Xoo lineages in Asia (Fig 4a; S8a-e Fig). Ancestral populations within the AXooL1 lineage appeared first around 6,427 years before present (yBP) (95% highest posterior density [HPD]=11,719-2,902 yBP) represented by AXoo1. Then, AXooL2 lineage exhibited by the node of AXoo5-7 surfaced around 4,080 yBP (95%HPD=8942-660 yBP). Finally, the AXooL3 recombinant group appeared around 808 yBP (95%HPD=1009-627 yBP). The AXoo1 divergence may have happened quite early, which could explain why AXoo1 was the most diverse and carried a high number of population-specific genes (Fig 1c, S5a Fig). Most of the AXoo1 isolates were collected from South China (Fig 2a), which are near the central Himalayan and Hengduan mountain river basins [20]. Previous reports have suggested that the wild rice populations from the same region might be the ancestors of cultivated *japonica* rice [11] or that they have feralized with cultivated rice in this region [56]. In either case, this might suggest that an ancestor of AXoo1 has adapted to a *japonica*-like population first. AXoo1 has *PthXo2* and *PthXo3* (Fig 4b; S5b Fig) TALEs capable of inducing *OsSweet13* and *OsSweet14*, respectively. This might indicate that *OsSweet13* and *OsSweet14* were targeted first. Interestingly, AXoo1 carries a version of *PthXo2* that activates the *japonica* allele of *OsSweet13* (Fig 4b).

A second group that might have emerged around ∼4000 yBP was AXooL2. AXoo7’s ancestor (∼3200 yBP) is a population that appeared in South Asia (Fig 4a; Fig 2a), where *indica* rice was domesticated [7], while the other populations within this lineage appeared at least less than 2000 yBP ago. Interestingly, AXoo7 carries a different version of *PthXo2* (Fig 4b; S5b Fig) that induces the *indica* alleles of *OsSweet13* [33,57]. The appearance of AXoo7 coincides with the first record of *PthXo1*, the effector activating *OsSweet11*, which is mostly distributed within AXooL2 members in South Asia (Fig 4b). Most AXooL2 populations in South Asia (AXoo6, AXoo7, and AXoo9) relied exclusively on *PthXo1* effectors (Fig 4b), suggesting that these populations evolved a mechanism to activate the third *OsSweet* target, *OsSweet11*. Therefore, it is not surprising that resistance gene *xa13*, a mutation in the promoter sequence that blocks the activation of *OsSweet11*, emerged in *indica* landraces [58].

Lastly, the appearance of AXooL3-derived populations ∼800 yBP (Fig 4) can be postulated between multiple recombination events between AXooL1 and AXooL2 (Fig 3c and S5c Fig), as rice is becoming a predominant crop in Asia. The first population to emerge from this lineage was AXoo11, which has the *PthXo2*-japonica version. This was followed by AXoo12, whose strains have either of the *PthXo2* versions. Finally, AXoo10 emerged, and it only has the *indica* type. It is tempting to speculate that the initial AXoo10 population could possibly be a japonica leaning pathogen that gradually could infect indica genotype through adaptation or an acquired novel virulence mechanism by recombination, and the same might be also applied to AXoo12. As previously described, AXoo12 strains could have almost all SWEET-targeting TALEs (Fig 4b; S6b Fig), and this might be because they could recombine easily with other populations and can adapt faster to new ecosystems, as observed with their rampant dispersal around Asia (Fig 2a).

In general, it is likely that AXoo lineages evolved alongside the domestication of rice. However, our results showed long HPD estimates that could bias the interpretation of the results, even though the population datasets used exhibited positive temporal signals befitting for molecular clock analysis. Increasing the sampling distribution to include countries absent in this study or adding AXoo infected rice herbarium samples collected at a certain time period in the past could potentially alleviate this problem.

### Asian Xoo could have followed the rice dispersal across Asia

Archeological and genetic evidence suggest that *japonica* and *indica* were domesticated independently and dispersed in distinct patterns [7,14]. To understand the migration pathway of AXoo, we performed an ancestral state reconstruction using Bayesian stochastic search variable selection analysis [59] and found that AXoo followed the rice dispersal pattern across Asia (Fig 5; S8a Fig). Predictions from BEAST showed that AXoo1 probably originated in China and could have dispersed west- and southward into South Asia and Southeast Asia (Fig 5). The timing of divergence (∼6,500 yBP) and the geographic origin might indicate that the progenitors of AXooL1 were in contact with a *japonica* lineage in this area. While *japonica* was domesticated in east China around 9,000 yBP [7,60], and archaeobotanical records indicate that *japonica* spread into South China between ∼5,000 yBP and ∼4,000 yBP. These values are supported by the demographic reconstruction showing a major relocation of *japonica* into South China between ∼4,500 to 4,000 yBP due to a cooling event, and further movement south towards Southeast Asia[14]. Although we can’t exactly pinpoint the exact migration pathway of AXooL1, we could hypothesize that AXooL1 emerged somewhere in China, and from there probably have followed japonica dispersal in Asia.

**Fig. 5.**
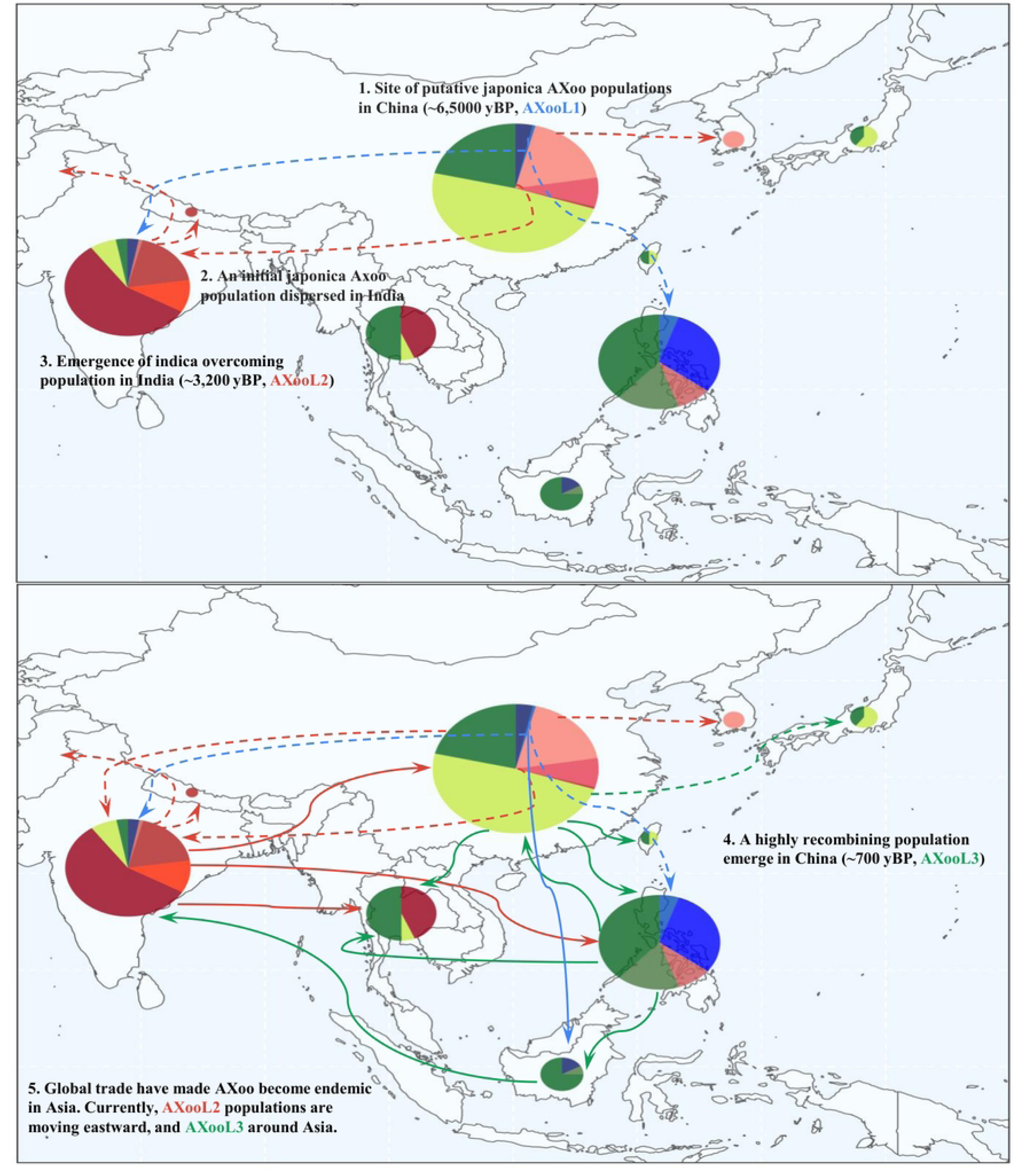
The hypothetical migration pathway of AXoo populations in Asia. The top panel visualizes the emergence and dispersal of AXoo populations > 1000 years before present (yBP), and the bottom panel shows < 1000 yBP. Initially, AXoo populations could have appeared in China ∼6500 yBP and evolved to cause disease in *O. sativa* subspecies *japonica*. The japonica AXoo populations must have moved to India, and probably along with japonica rice varieties as. Then, *O. sativa* subspecies *indica* domestication happened in India, and the existing japonica AXoo populations were essentially the seed population for the emergence of indica overcoming AXoo around 3200 yBP. Then, due to the genetic mixing of different AXoo populations that are compatible with different rice varieties, and also with the initial global trade at that time i.e the Silk Road, a third highly recombing AXoo population might have emerged in China at least 700 yBP. Since then, AXoo populations have been circulating around major rice growing countries in Asia. The migration lines are based on the ancestral state prediction performed in BEAST. The solid lines denote the certainty of the prediction, and while the dotted lines show uncertainty of the prediction due the absence of a temporal signal or data. The color of each line represents AXoo lineages: blue = AXooL1, red = AXooL2 and green AXooL3. The pie chart shows the proportion of the population in each country, and the size pertains to the number of sampled strains.

The demographic prediction placed the origin of the indica overcoming AXooL2 in India (Fig 5). The oldest AXooL2 population (AXoo7) emerged around 3,000 yBP (Fig 4a), which matches with the domestication of *indica* rice in the Ganges River basin around 4,000 yBP [14]. Although the timing of AXoo4 could not be predicted, its evolutionary history can be linked with the *japonica* adapted AXoo populations in South China since the population carries the japonica verison of *PthXo2* (Fig 4b) and dispersed to India, where it adapted to *indica*. AXoo4 shares the same node with the rest of the AXooL2 lineage (Fig 1a), and strong gene flow was observed with AXoo7 (S5c Fig). AXooL2 then moved westward from India into China (Fig 5), and this migration could explain the recent reintroduction of AXooL2 population (AXoo5).

The BEAST analysis also suggested that AXooL3 emerged in China (Fig 5), ∼800 yBP, after one or multiple recombination events (Fig 3). The actual distribution pattern and the timing can only be explained by increasing human activity. The Silk Road which flourished during the thirteenth and fifteenth centuries was the main sea and land trade pathway and usually involved key commodities such as rice, beans, and cotton [61]. This endeavor might have spread members of this lineage to almost all Asian geographies. While it is unclear why only AXooL3 members might survive the journey, other dispersal hypotheses are equally plausible. In any case, it appears likely that the history of AXoo is linked to the dispersal pattern of rice in Asia.

Due to global trade, AXoo has become more endemic in Asia. AXooL2 has recently moved eastward from India to Southeast Asia (Fig 5), as exemplified by a *xa5* resistance gene-breaking strains [26].Conversely, AXooL3 has rapidly spread across various Asian regions (Fig 5), and an example of which is the emergence of AXoo12 in the Philippines (S2a Fig) that could break *Xa4* resistance during the “Green Revolution” [22]. Moreover, a recent report revealed the presence of AXoo phylogroup in East Africa for the first time, threatening rice production there [62]. Rice resistance breeding, either through conventional methods or the use of modern techniques such as genome editing [33], has been shown to be successful in controlling the AXoo pathogen. However, it is necessary to be informed of the local pathogen populations of a country or region first to properly implement precise disease management strategies.

## MATERIALS AND METHODS

### *Xanthomonas oryzae pv. oryzae* dataset collection and sequencing

A total of 433 Asian Xoo genome sequences were used (S1 Dataset) and were obtained from public databases (421) and from this study (12). The 421 AXoo genomes dataset were compiled from published reports in India [19], the Philippines [22], Thailand [26,63], China [20] and from other countries [33]. To sequence new strains from Indonesia, Xoo cultures of twelve isolates were grown overnight at 30 °C, and extraction of DNA was done using the Easy-DNA kit (Invitrogen, USA) following the manufacturer’s protocol. Library preparation and pair-end sequencing using Illumina HiSeqTM 4000 were performed by Beijing Genomic Institute (BGI, Shenzhen, China). The filtering of the raw reads which includes removing adapter sequences, contamination and low-quality reads were conducted by BGI.

To compile the phenotypic data, the reported data [20,22,63,64] of 344 Xoo strains over a set of seven near-isogenic lines carrying single resistance genes were collected: IRBB4 (Xa4), IRBB5 (xa5), IRBB7 (Xa7), IRBB10 (Xa10), IRBB13 (xa13), IRBB14 (Xa14), and IRBB21 (Xa21). In order to consolidate the phenotypic reactions and reduce variation of the quantitative of the data, since all the source papers have different cutoff for determining Resistant (R) or Susceptible (S) and the measuring time of disease observation (days post inoculation (dpi)), we scaled them into two categories, R < 10 cm and S > 10 cm. The new categories were applied for only three papers. For Mishra et al. [64], observations were taken 14 dpi and lesion lengths were measured R < 5 cm, 5-10 cm = moderate and S > 10 cm, and the new scale for this dataset are described above. The same case was applied for the Quibod et al. [22] dataset, as observations were captured 14 dpi and lesion length scales were designated as <5 cm = resistant (R), 5–10 cm = moderately resistant (MR), 10–15 cm = moderately susceptible (MS), and >15 cm = susceptible (S), and was converted into R and S only as mentioned above. In the case of the phenotypic data from Song et al. [20] which measured the lesion length 21 dpi, we converted the quantitative lesion length data into qualitative following the new designation described above. The only exception was the data from Chintaganon et al. [63], where S was considered a lesion length ≥ 6 cm and R at < 6 cm.

### Genome assembly and re-annotation

De-novo assembly was performed on all 12 newly sequenced AXoo isolates. Reads from Illumina were assembled into contigs and scaffolds using SPades v3.11.1 [65]. The following parameters were applied: -careful (to reduce mismatches and short indels), --cov-cutoff auto (to allow the program to find the optimal coverage cutoff), and -k 21,33,55,77 (k-mer sizes). All the assembled genomes were checked for quality using CheckM v1.2.2 [66]. Gene-calling and annotation were done to all draft and complete Xoo genomes using Prokka [67]. The compiled GenBank files of AXoo genomes used in Oliva et al. [33] served as the primary annotation source.

### SNP calling, phylogenetic reconstruction, and population structure analysis

Harvest suite 1.1.2 [68] which houses Parsnp, a rapid multiple genome alignment tool was applied to obtain the core genome of all 433 strains, and extract single nucleotide variants from the core alignment. The core genome consisted of a ∼2.3 Mb with ∼21,900 SNPs. We detected loci that were under recombination using ClonalFrameML v1.12 [41] with 100 simulations to approximate uncertainty. The recombining regions were then masked (https://github.com/kwongj/maskrc-svg) and subjected to phylogenetic analysis. A maximum likelihood (ML) phylogeny was ascertained with RAXML 8.2.12 [69], and phylogenetic node statistical confidences were inferred from 1000 bootstrap runs using the general time reversal with the Gamma model of rate heterogeneity as a nucleotide substitution model. The ML phylogenetic tree was visualized with the R package *ggtree* [70]. We also employed the same pipeline to further characterize the identified lineages, populations with specific reference genoms, and with the *Xanthomonas oryzae* pv. *oryzicola* BLS256 (Accession Number: CP003057.2).

The SNP dataset from 433 genomes was used to predict the population structure. Initial population clustering was predicted using the R package *rheirbaps* [71] which implements the hierarchical structuring algorithm from the BAPS program. Two independent runs were performed resulting in two levels of nested genetic populations: 15 (1^st^ level) and 51 (2^nd^ level). The inferred population structure was further constricted into twelve main groups by fastBAPS optimized symmetric algorithm [23], incorporating both the SNP calls and the phylogenetic tree. Additional analyses, including evolutionary distance estimation [72] and a pairwise SNP distance matrix (https://github.com/tseemann/snp-dists), were conducted to support the twelve clusters. The results of these supplemental analyses were used for principal component analysis (PCA).

### Pan-genome analysis and prediction of potential pathogenicity genes

The pangenome of the 433 AXoo genomes was constructed using PEPPAN v1.0.5 [73]. A total of 7,585 orthologous gene groups (S3 Dataset) were clustered and grouped into four categories: hardcore (99% <= strains < 100%, n = 3,067), softcore (95% <= strains < 99%, n = 290), shell (15% <= strains < 95%, n = 1393) and cloud (strains < 15%, n = 2835). The core genes were aligned with MAFFT v7.453.0 [74] with the FFT-NS-i alignment method. Furthermore, the core genes were used to calculate nucleotide diversity (pi), Wright’s fixation index (F_ST_), and Tajima’s D using the R package *PopGenome* [75] (S4 Dataset). Diversity analysis was executed within and between population and within countries. The orthologous groups were searched for potential pathogenicity signatures. Using blastn [76], candidate Xanthomonas outer proteins (*Xop*) were identified using the Xanthomonas database (http://www.xanthomonas.org/) with at least 70% identity and 100% coverage. Genes involved in signaling and regulation of virulence were also examined by blastn against a comprehensive database [77]. Carbohydrate-active enzymes (CAZymes) with signal peptides that are candidate cell wall degrading enzymes (CWDE) were identified using the automated annotation pipeline in dbCAN2 [78]. To identify putative secreted proteins that might be involved in pathogenicity, secretion signals were predicted with SignalP 5.0 [79], and inferred for the presence of transmembrane domain-containing proteins was predicted using TMHMM 2.0 [80] and PRED-TMBB2 [81]. Proteins with transmembrane domains were removed, leaving a final list of proteins inferred to be putative secreted proteins that might be involved in pathogenicity.

### Distribution of transcription activator-like effectors (TALE) in AXoo

Completely assembled level AXoo genomes were subjected to TALE annotation. A total of 41 genomes that were sequenced using long-reads were used to identify the TALE positions using the program AnnoTALE v1.4.1 [82]. For the TALE phylogeny, nucleotide TALE sequences served as input for DisTAL v1.1[83], which computes distances from the TALE repeats, and builds a neighbor-joining tree. The phylogenetic tree was visualized using *ggtree*. The TALE classification was based on a previous grouping [18] and updated for newer clusters (S2 Datatset). An RVD-based string clustering was performed using Damerau-Levenshtein distance with at least eight string differences implemented in the R package clustringr (https://github.com/cran/clustringr) to support the allele sharing of each TALE group. In addition, TALE classification from the AnnoTALE database was also added.

### Detection of recombination

Three methods were used to detect the presence of recombination in AXoo. The initial assessment was performed using the Neighbor Net method implemented in SplitsTree4 v4.16.1 [40] to observe recombination patterns within the pathovar, and utilized a pairwise homoplasy index (PHI) to test for the presence of recombination. Single nucleotide variants were used as input to visualize the phylogenetic network. Recombination segments detected within the core genome alignment from parsnp were supplied to ClonalFrameML v1.12 [41] and the orthologous gene groups for fastGEAR [42]. Recombination prediction from fastGEAR was conducted using a predefined population partition and default specification on the gene alignment, and the results are shown in S3 Dataset. Recombination events were then categorized as recent (recombination affecting a few strains) and ancestral (recombination affecting the entire population) as described by fastGEAR. The software could detect the possible origin of recombination, so we classified the flow of the recombination events as either donor (source of the recombination segment) or recipient (receiver of the recombination fragment).

### Bayesian phylogenetic inference of the Xoo population

For creating the bayesian molecular clock trees, an initial analysis to test the presence of a significant positive correlation of our heterochronous data with ML phylogeny was performed using root-to-tip regression and random permutation of sampling dates. SNP calling and ML tree construction were performed for each AXoo population independently or in combination with other AXoo populations of the same lineage and for the whole AXoo lineage. Next, the temporal signal of the isolates with heterochronous data was tested using TempEst v1.5.3 [84]. We further checked the appropriateness of our temporal datasets by performing a random permutation of sampling dates [85], with 1,000 permutations for each combination. From the results, using all of the AXoo genomes didn’t yield a positive correlation between time and root-to-tip divergence in the regression analysis (S7a Fig). However, we observed clock-like behavior for lineage AXooL3, combining AXoo5, 6 and 7, and individual populations of AXoo1, AXoo8 and 9 (S7b-c Fig). BEAST v 1.10.4 [48] was used to execute the temporal Bayesian phylogenetic analysis to estimate the timeline of the most recent common ancestor (MRCA), the evolutionary rate, and the phylogeographic dynamics. A mix of three clocks (strict, relaxed log-normal, and relaxed exponential) and three population dynamic (constant, exponential, and skyline) models were applied. For each model combination, the run was allowed to continue for 300 million Markov chain Monte Carlo (MCMC) and sampling the posterior every 1000th iteration with general time-reversible (GTR) as the nucleotide substitution parameter. Model combinations that failed to converge and or had less than 200 effective sampling sizes (ESS) were not considered. The clock rates and population models with the highest Bayes factor were chosen as the final representative of the analysis. We repeated each run three times and combined all the logs and trees, after considering 10% of the chain as burn-in using LogCombiner v1.10.4. The maximum clade credibility (MCC) tree was summarized in TreeAnnotator v1.10.4 and visualized through the R package *ggtree*.

For inferring the phylogeography dynamics, the ancestral state reconstruction for each lineage or population was predicted using a discrete trait evolution model [59]. Bayesian stochastic search variable selection analysis was turned on and a symmetric substitution model was used for reconstructing the ancestral state. The variable used for the discrete model was the country of origin of each isolate.

### Data Availability

The sequences produced in this study are available under NCBI BioProject PRJNA1031561. Additional datasets are under Dataset S1-4. Annotated genomes, scripts and genomic analysis are available at https://zenodo.org/records/14749993.

## Acknowledgment

Scientists at IRRI are funded by the Initiatives (PHI, ABI) of the Consortium for International Agricultural Research (CGIAR) and other bilateral projects funded by the Ministry of Agriculture, Taiwan; National Insititute of Crop Science (NICS), Korea; Commonwealth Scientific and Industrial Research Organisation, Australia. MHN was supported by a scholarship from the University of Southern Queensland. We thank the Philippines DOST–Advanced Science and Technology Institute (DOST-ASTI) for free access to high-performance computing. This work was funded by the Deutsche Gesellschaft für Internationale Zusammenarbeit (GIZ) through the Fund International Agricultural Research (FIA), grant number: 81219435 and by the Bill & Melinda Gates Foundation (Investment F-2016/1166-4, INV 008226, and OPP1088843). ILQ is supported by the ANR project PHRACE (ANR-22-CE20-0016).

## SUPPORTING INFORMATION

### Supplementary Figures

**S1 Fig. Phylogenetic construction of Asian Xoo (AXoo) reveals three main lineages.** An unrooted maximum-likelihood tree of Asian Xoo with *Xanthomonas oryzae* pv. *oryzicola* (Xoc) as an outgroup.

**S2 Fig. Population structure analysis reveals twelve *Xanthomonas oryzae* pv. *oryzae* populations in Asia (AXoo)** a) A maximum-likelihood tree was constructed using 22,115 non-recombining SNPs. The predicted population structures reported in the Philippines (PX-a, PX-B, PX-C), India (L-I, L-II, L-III, L-IV, L-V), and China (CX-1, CX-2, CX-3, CX-4, CX-5, CX-6) are shown [1, 2, 5]. The new designated Xoo populations (AXoo1-12) were inferred using fastBAPS [18]. The red dots in the node of the phylogeny are bootstrap values greater than 95. The population structure from the previous analysis coincides with the predicted groupings done in this study. b) Principal component analysis of the pairwise distance matrices of 433 assembled genomes showed clustering of 12 populations. An alignment-free method [19] was used to acquire pairwise evolutionary distances between closely related genomes. c) Principal component analysis of the pairwise distance matrices using SNPs. snp-dist was used to convert SNP fasta alignments into pairwise SNP distance matrices.

**S3 Fig. Temporal distribution of *Xanthomonas oryzae* pv. *oryzae* populations in Asia (AXoo).** The temporal distribution of each AXoo population in the form of treemaps. The boxes are arranged and sized according to the total number of input isolates in each given population and timeframe.

**S4 Fig. Pan-genome landscape of *Xanthomonas oryzae* pv. *oryzae*.** a) The pan-genome consists of 7,585 orthologous genes categories as hardcore (99% <= strains < 100%, n = 3,067), softcore (95% <= strains < 99%, n = 290), and accessories with shell (15% <= strains < 95%, n = 1,393), and cloud (strains < 15%, n = 2,835). b) The rarefaction curve shows Xoo has a close pan-genome. The plot illustrates the core (solid line) and accessories (dashed line) gene numbers in the twelve AXoo populations. Only AXoo9 follows an open pattern of pan-genome. The median values from 1,000 random permutations of ortholog groups from the 433 genomes were collected to visualize the rarefaction curves. c) Heatmap of the number of shared core genes among AXoo populations. Most of the populations within the lineage share the same alleles. d) Heatmap of the number of shared accessory genes among AXoo populations. Most of the populations within the lineage share the same alleles.

**S5 Fig. Patterns of diversity and population differentiation across *Xanthomonas oryzae* pv. *oryzae* populations in Asia (AXoo)** a) Nucleotide diversity (*Pi*) of AXoo populations using the core gene sequences. AXoo1 was the most diverse at an average of 0.98445863, followed by AXoo7 (0.75185412) and AXoo12 (0.49216247). The least diverse population was AXoo5 (0.05750352). b) Tajima’s D (*D*) computed for the AXoo population. The majority of AXoos have negative median values for Tajima D except for AXoo8 with a positive value. c) The heatmap displays the mean Fixation index (*Fst*) values. Populations within lineage AXooL3 have the lowest *Fst* values suggesting continuous gene flow within these populations. Additionally, AXoo1 and AXoo7 have decreasing patterns of *Fst* across populations which could imply that both populations are the source of the genetic diversity.

S6 Fig. Distribution of Transcription activator-like (TAL) effector on Asian populations of *Xanthomonas oryzae* pv. *oryzae* (AXoo). a) Neighbor-joining tree built using the distance matrix from the program DisTAL [29] based on alignments of repeat regions from TAL effectors. A total of 24 predicted TAL effector families (TEF) were identified based on TEF families [38]. b) Each population harbors different RVD alleles of the TAL effectors. AXoo1 has fewer TEF compared to other populations. AXoo10, AXoo11, and AXoo12 contain the most alleles. The first five TEF are considered as the core, while the rest are accessories. The number of representative genomes per each population is indicated. TAL effector alleles from 1 to 5 are represented by more than or equal to two representative isolates, while “other” denotes a singleton allele. No long read sequences were available for strains belonging to the AXoo5 population.

**S7 Fig. Testing for molecular clock signals in different *Xanthomonas oryzae* pv. *oryzae* group combination.** The correlation between time and root-to-tip divergence was analyzed a) for all of the 375 genomes with temporal data, b) for each of the three lineages and combining populations AXoo5, 6, and 7, and c) for each population. Groups with possible temporal signals are highlighted in blue. The black solid line represents the regression line, and the dashed lines show the 95% confidence interval. Regression coefficient was used to estimate the fit of the data, and the p-value that compares the true R-value to the R-value estimated from 1000 random permutations.

**S8 Fig. Temporal phylogenetic reconstruction of lineages and population with appropriate clock signal using BEAST** [36]. Maximum clade credibility trees were presented for a) AXoo1 (AXooL1), b) AXoo5-7 (AXooL2), c) AXoo8 (AXooL2), d) AXoo9 (AXooL2), and AXoo10-12 (AXooL3). The branch color corresponds to the country with greater support. Data along the x-axis are in calendar years (CE). The black bars in each node are error bars showing the 95% highest posterior density (HPD), and while the colored points inside the nodes denote >=90 posterior probability and th with the highest posterior probability of ancestral states.

### Supplementary Table

**S1 Table. Molecular clock estimated time since most recent common ancestor (TMRCA) and nucleotide substitution rate per site per year.** The TMRCA and substitution rates were inferred using BEAST v 1.10.4. [36].

### Supplementary Dataset

**S1 Dataset. General information of all *Xanthomonas oryzae* pv. *oryzae* (Xoo) strains used in this study.** The columns are partitioned into different categories. The columns Asian Region to Longitude are data relating to geography. The labels under Asian Region denote the following: EA (East Asia), SEA (Southeast Asia), and SA (South Asia). The columns from IR24 to IRBB21 are pathotyping assays of near-isogenic lines containing single bacterial blight Xa genes. The columns Lineage to *rheirbaps* shows the different sources of the population structure analysis. The population analyses that were carried out from previous studies are columns Philippines Population [2], China Population [5] and India Population [1]. The citations in the reference column are listed below.

S2 Dataset. The list of Transcription Activator-Like Effectors (TALE) annotated from 41 complete genomes of *Xanthomonas oryzae* pv. *oryzae* strains. The column TALE-annotations are CDS predictions obtained from the program AnnoTALE [28]. The classification of TALEs is in the columns TEF (TALE families) classification [38], AnnoTALE Class, and RVD String Clustering (https://github.com/cran/clustringr). The asterisks in the column TEF classification/Known TALE are predicted to be new families in the TEF classification. Alleles in each TALEs are described in the last column (TALE Alleles). Another thing to note, AnnoTALE prediction annotated iTALEs as pseudo TALEs.

**S3 Dataset. The list of orthologous gene groups under recombination from the fastGEAR result.** The table also shows the pangenome category of the orthologous group.

**S4 Dataset. Genetic diversity values for each gene either within and between populations.** The values were computed using the R package *PopGenome* [33]. The results presented in the sheet are as follows: nucleotide diversity (Pi), Tajima’s D and Fixation index (Fst).

